# Bifunctional Lipid-Protein Crosslinking Efficiency and Reaction Products

**DOI:** 10.64898/2026.01.18.700185

**Authors:** Carla Kirschbaum, H. Mathilda Lennartz, Katelyn C. Cook, Kristin Böhlig, Athanasios Papangelis, Carol V. Robinson, André Nadler

## Abstract

Bifunctional diazirine lipids are valuable tools for mapping protein–lipid interactions and cellular localization by photocrosslinking. Yet, the crosslinking efficiency of these probes has not been systematically evaluated. Here, we use the lipid transfer protein STARD10, which binds phospholipids in a 1:1 stoichiometry within a hydrophobic pocket, to measure the upper limit of the photo-crosslinking efficiency of bifunctional lipid probes. We characterize reaction products using native and denaturing mass spectrometry. Our results show that approximately 5% of photoactivated lipids form covalent protein–lipid crosslinks, while the majority follow intramolecular reaction trajectories, resulting in the formation of products featuring alkene, ketone and hydroxyl moieties. These findings provide essential context for the use of bifunctional probes to uncover the cell biology of lipids and highlight the need for continuous improvement to experimental workflows.

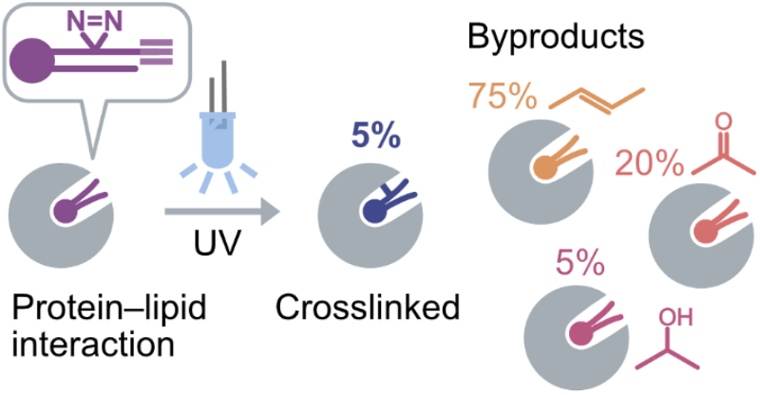

---

Diazirine photoreactive groups are widely used to investigate protein–protein and protein–small molecule interactions by photoaffinity labeling.^1–5^ Diazirine-containing phospholipids closely mimic the behavior of natural lipids and readily mix with endogenous membranes, making them particularly valuable for probing protein–lipid interactions.^6, 7^ Bifunctional lipids that combine a diazirine moiety with an alkyne tag have emerged as powerful probes for mapping lipid interactomes^8–16^ and are currently the most reliable tools for monitoring species-specific intracellular lipid localization and transport at high resolution.^17–20^

The reactivity of dialkyl diazirines has become increasingly well understood in recent years. Upon UV-A irradiation, they generate highly reactive carbenes through the expulsion of N_2_, but also form substantial amounts of diazo intermediates (*cf*. **Figure 1A**).^21, 22^ Those can either convert into carbenes or react directly with proteins.^21^ Unlike carbenes, diazo intermediates have a chemical lifetime above the diffusion limit^23^ and react over a wider radius, with a high selectivity for acidic and polar amino acids.^24–27^ Together, the rapid, diffusion-limited carbene pathway and the slower diazo-mediated pathway shape the final product distribution.^22^

**Figure 1.**
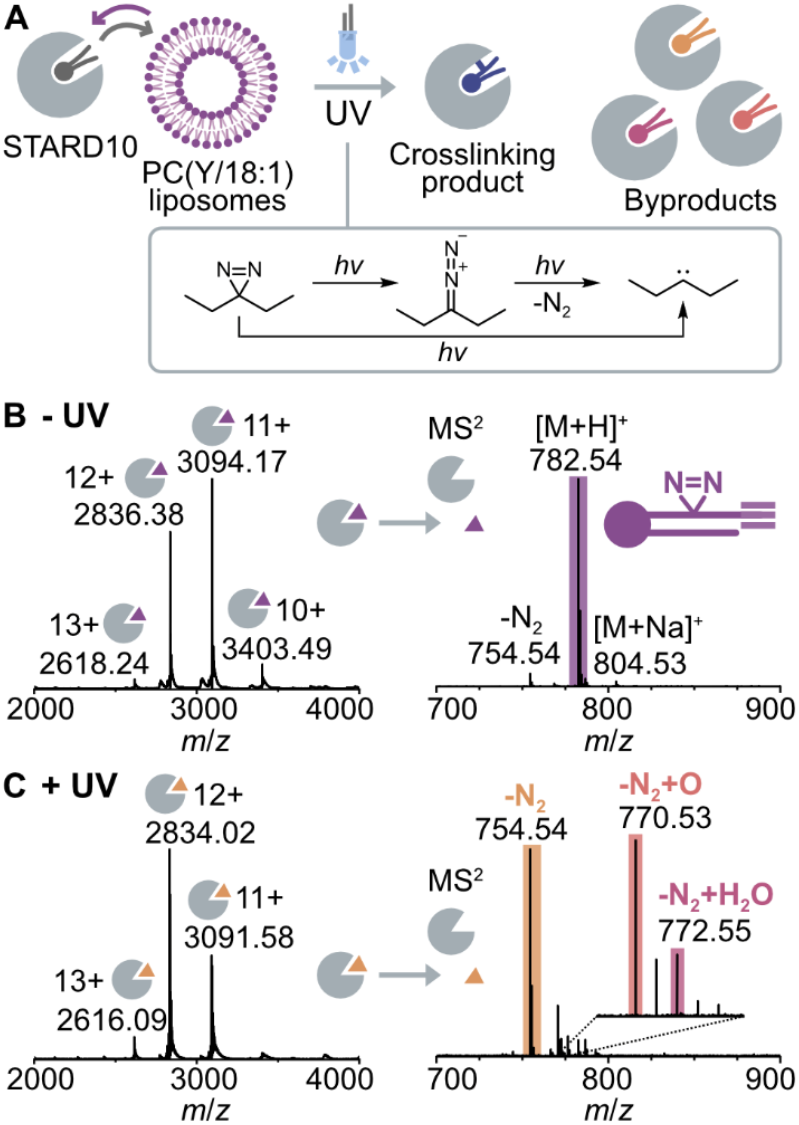
Crosslinking of STARD10 and bifunctional PC *in vitro*. (A) STARD10 is incubated with liposomes containing 80 % PC(Y/18:1) and 20 % cholesterol to replace copurified bacterial lipids (dark gray) with diazirine PC (purple). Diazirine PC forms reactive diazo and carbene intermediates upon UV irradiation. (B) Native MS before photoactivation confirms saturation of the 1:1 STARD10:PC(Y/18:1) complex. Colored triangles represent bound lipids. (C) Native MS of STARD10 and release of non-crosslinked lipids after UV irradiation shows nitrogen loss products and a minor fraction (ca. 25 %) of oxidized lipids.

The efficiency of the protein–lipid crosslinking reaction remains undefined, though it is one of the key experimental constraints when studying lipid localization and lipid-protein interactions. Model peptides have shown crosslinking rates of up to 100% when exposed to excess diazirine reagent, but these rates drop to nearly zero in aqueous solution.^21^ In 2:1 phosphatidylcholine (PC):diazirine-PC liposomes containing model transmembrane peptides, ca. 1–10% peptide–lipid crosslinking products were recovered after repeated irradiation,^6, 7^ but those values are highly system-dependent. It is well-established that the crosslinking efficiency is significantly reduced by intra- and intermolecular side reactions of the reactive intermediates. Notably, intramolecular alkene formation of photoactivated diazirines can account for >50% of the reaction products,^22^ and lipid–lipid crosslinking is frequently observed in liposomes.^6, 7^ We sought to determine the upper limit of lipid–protein crosslinking efficiency and the extent of the competing intramolecular reaction pathways, while reducing side-reactions with other lipids and small molecules.

To this end, we used the soluble lipid transfer protein STARD10 and bifunctional phosphatidylcholine (PC) derivatives as a model system. STARD10 binds PC in a 1:1 stoichiometry within a hydrophobic binding pocket that excludes water.^28^ Hence, any lipid molecule activated inside this pocket can, in principle, undergo only two possible reactions: crosslinking to the protein or internal rearrangement. As STARD10 forms a 1:1 complex with the respective lipid probe, the relative fraction of covalent protein–lipid conjugates and rearranged lipid products can be directly determined under ideal, solvent-free conditions.

We used native and denaturing mass spectrometry (MS) to characterize the reaction products and quantify the crosslinking efficiency after a 3-second irradiation pulse with 365 nm high-power LEDs. For STARD10 saturated with bifunctional PC, we observed a crosslinking yield of approximately 5%, whereas the majority of bound lipids formed alkenes, and a minor fraction resulted in oxidation products. Our results suggest that a significant loss of signal-to-noise ratio during the photocrosslinking step should be taken into account when planning experiments with bi-functional lipids.

We obtained human STARD10 by recombinant expression in *E. coli*. The protein copurified with bacterial phospholipids in a 1:1 stoichiometry, as confirmed by native MS (**Figure S1**). To replace the natural phospholipids with diazirine lipids, we incubated purified STARD10 with liposomes consisting of 80 % bifunctional PC(Y/18:1) (full structure shown in **Figure 3A**) and 20 % cholesterol in PBS solution (**Figure 1A**). After 10 min incubation at room temperature, we removed excess lipids by buffer exchange into ammonium acetate (200 mM, pH 7) for native MS.

The native mass spectrum of STARD10 incubated with PC(Y/18:1) showed a charge state distribution from 10–13+ corresponding by mass to STARD10 bound to PC(Y/18:1) in a 1:1 stoichiometry (**Figure 1B**). To confirm the identity of bound lipids, we isolated the protein–lipid complex in the ion trap and used collisional activation to release non-covalently bound lipids from STARD10. The MS^2^ spectrum confirmed that the bound lipids corresponded to intact PC(Y/18:1). A minor peak indicated partial nitrogen loss, which we ascribe to the collisional activation. Overall, the MS data confirmed that the diazirine lipids had entirely replaced the bacterial phospholipids copurified with STARD10, providing ideal conditions for assaying crosslinking efficiency and reaction products.

We next irradiated the 1:1 STARD10/PC(Y/18:1) complex for 3 s using a high-powered 365 nm LED. After photo-activation, we removed excess lipids by buffer exchange and investigated the reaction products by native MS. Consistent with the expected loss of the diazirine group upon UV photoactivation, we observed a decrease in the mass of the protein–lipid complexes relative to the control without UV irradiation (**Figure 1C**). To characterize the reaction products, we isolated the protein–lipid complex in the gas phase and released non-covalently bound lipids by collisional activation. The majority of reaction products (ca. 75 %) corresponded by mass to the loss of N_2_ (−28 Da), whereas additional peaks shifted by +16 and +18 Da suggested partial lipid oxidation (ca. 25 %). The precursor made up only 5 % of the bound lipids after photoactivation, suggesting that it was almost entirely consumed within the 3-second irradiation pulse.

Non-covalent protein–lipid complexes are indistinguishable by mass from covalent protein–lipid crosslinking products (**Figure 2A**), rendering native MS unsuitable for determining the fraction of STARD10 that was efficiently cross-linked. To disrupt non-covalent protein–lipid interactions while preserving covalently bound lipids, we denatured the protein in organic solvent (45% acetonitrile, 5% isopropanol, 1% formic acid). Denaturing MS of STARD10 incubated with PC(Y/18:1) in the absence of UV irradiation yielded a broad charge state distribution from 10–41+ corresponding to the unmodified apo protein (**Figure 2B**). Deconvolution of the spectrum confirmed that only negligible amounts of lipid remained associated with the protein after denaturation (ca. 1 %), demonstrating effective removal of non-crosslinked lipids.

**Figure 2.**
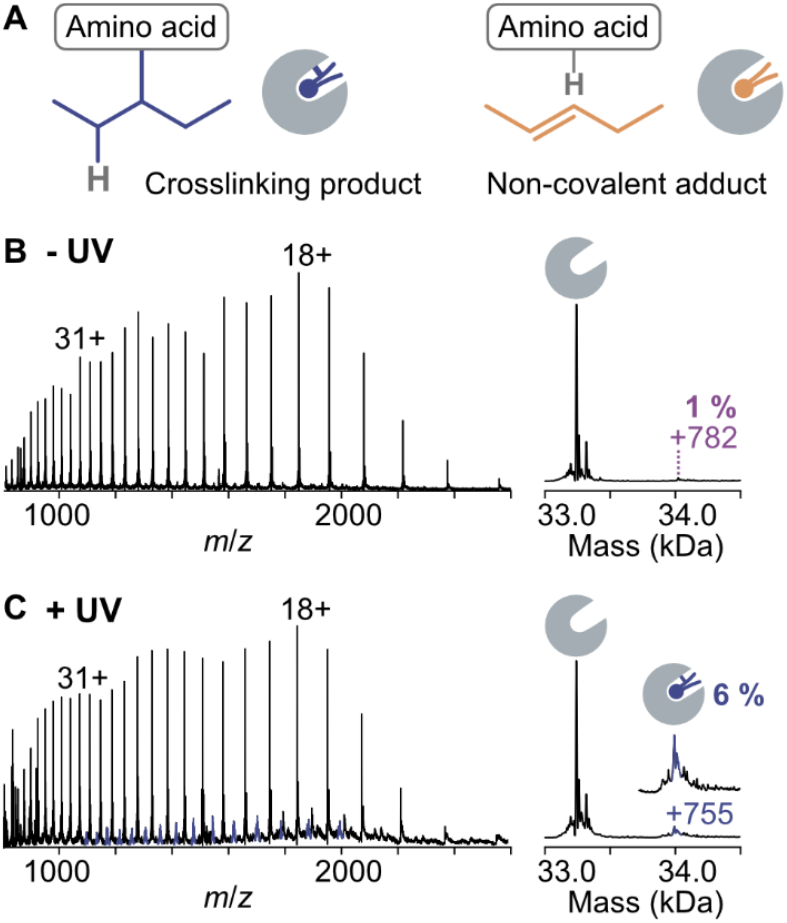
Crosslinking efficiency determined by denaturing MS. (A) Covalent protein–lipid complexes are indistinguishable by mass from non-covalently bound alkene byproducts. (B) Denaturing MS of STARD10 incubated with PC(Y/18:1) without UV irradiation confirms disruption of non-covalent protein–lipid interaction. (C) Denaturing MS of STARD10 incubated with PC(Y/18:1) and crosslinked with UV light reveals that photoactivation led to crosslinking in ca. 5 % of the cases, assuming 1 % residual non-covalent binding.

**Figure 3.**
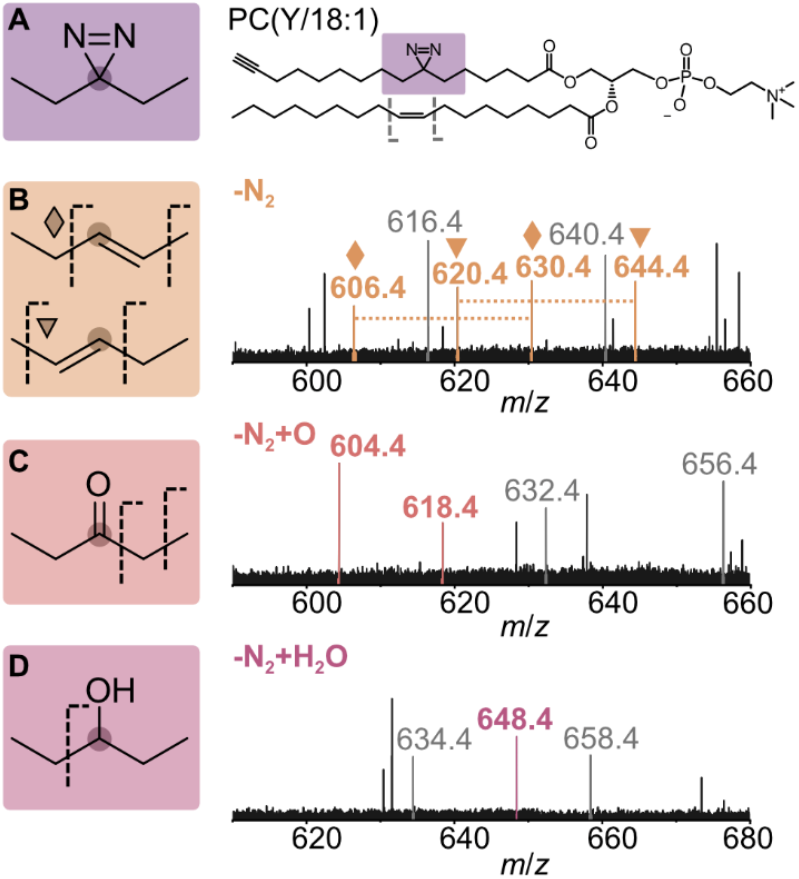
Characterization of byproducts by ultraviolet photo-dissociation mass spectrometry. Photoactivation of the diazirine lipid PC(Y/18:1) (A) bound to STARD10 resulted in the formation of two regioisomeric alkenes (B), ketones (C) and alcohols (D) inserted at the original position of the diazirine moiety. Dotted lines indicate C-C bond cleavage by UVPD. Peaks labelled in gray are derived from the C=C bond in 18:1 acyl chain.

To quantify the protein–lipid crosslinking efficiency, we performed denaturing MS of STARD10 incubated with PC(Y/18:1) and irradiated with UV light (**Figure 2C**). Besides the major charge state distribution corresponding to the unmodified apo protein, the mass spectrum featured a minor charge state distribution corresponding to STARD10 covalently bound to one PC molecule. We deconvoluted the spectrum using UniDec^29^ and determined by peak integration that ca. 6 % of the protein was crosslinked to PC at a 1:1 ratio. When accounting for 1 % of nonspecific lipid binding, the estimated crosslinking efficiency amounts to approximately 5 %. This ratio was consistent across technical replicates (**Figure S2**). Because the protein was saturated with PC(Y/18:1) prior to UV irradiation and the precursor was almost entirely consumed after irradiation, we conclude that ca. 5 % of the photoactivated lipid resulted in effective crosslinking, whereas the majority (approximately 70 %) led to nitrogen loss and intramolecular rearrangement without crosslinking to an amino acid.

To consolidate these values, we also quantified the cross-linking efficiency of bifunctional PC probes with alternative side chains (PC(Y/16:0) and PC(Y/20:4)), which could influence how the lipid chains are folded inside the STARD10 lipid-binding pocket (**Figure S3**). Like PC(Y/18:1), both lipids were readily loaded onto STARD10 via liposomes, though PC(Y/16:0) replaced the bacterial lipids less effectively than the unsaturated lipid probes. Denaturing MS revealed similar crosslinking rates between 5–7 % for PC(Y/16:0) and PC(Y/20:4), suggesting high reproducibility of the photoreaction pathways.

To further characterize the byproducts formed upon photoactivation that do not lead to covalent protein–lipid cross-linking, we measured ultraviolet photodissociation (UVPD) mass spectra of all reaction products (**Figure 3**). UVPD is an ion activation technique that induces cleavage of C-C bonds and thus yields deeper insight into lipid structure than classical collision-based fragmentation.

In the UVPD spectrum of the main byproduct corresponding by mass to the loss of N_2_, we detected two pairs of peaks shifted by 24 Da, which is indicative of alkene formation (**Figure 3B**).^30^ The fragment spectrum confirmed the formation of two alkene regioisomers with double bonds to the left and right carbon neighboring the original diazirine site. Note that the configuration (*E*/*Z*) of the newly formed C=C bond cannot be determined with this approach. The C=C bond in the native 18:1 acyl chain of the PC(Y/18:1) probe yielded an additional pair of peaks shifted by 24 Da (shown in gray). Based on the relative intensities of the fragment ions, we suggest that both alkene regioisomers are generated in comparable amounts.

The oxidized lipid products correspond most likely to a ketone (+16 Da) (**Figure 3C**) and an alcohol (+18 Da) (**Figure 3D**) inserted at the original diazirine site. For the ketone, we observed a fragment resulting from cleavage between the alpha and beta carbon, a reaction that has previously been observed for small molecules containing ketones.^31^ A minor fragment was also formed by C-C bond cleavage adjacent to the carbonyl oxygen. The alcohol yielded a diagnostic fragment that contained the newly formed OH group by cleavage of the C-C bond adjacent to the original diazirine site. Full fragment ion assignments supported by high-resolution Orbitrap mass spectra can be found in **Figure S4** and **Table S1**. Overall, we observed ca. 75 % alkene, 20 % ketone and 5 % alcohol formation in the original diazirine acyl chain.

The byproducts identified by MS are consistent with previous studies. It is known that alkene formation is the major side reaction of photoactivated dialkyl diazirines in liposomes and non-aqueous solutions.^7, 22^ Under aqueous conditions (80% water), photoactivation of a soluble dialkyl diazirine yielded a mixture of 50% alkene, 10-20% ketone and 35% alcohol through water insertion.^27^ The reduced abundance of oxidized lipids observed in our experiment is reasonable because the diazirine lipids are integrated into the hydrophobic binding pocket of STARD10, minimizing their exposure to water.

The average crosslinking efficiency of 5 % determined in this study is within the same order of magnitude as previous studies reporting 1–10 % crosslinking efficiency using 300 or 365 nm irradiation^6, 7^, although these values cannot be directly compared due to varying experimental conditions such as light source, irradiation time, and initial protein and diazirine lipid concentrations. Our results imply that 5% crosslinking yield represent a practical upper limit that can be achieved in the case of a well-defined 1:1 protein:lipid complex featuring a tightly bound lipid within a geometrically confined binding pocket. This allows for the more general conclusion that a large majority of diazirine phospholipids will not form crosslinks to proteins in cellular assays. A larger number of protein–lipid crosslinks is expected statistically for proteins that interact with multiple lipids, such as membrane proteins which are preferentially labeled by dialkyl diazirine probes,^27^ though the intrinsic chemistry of each individual lipid probe remains unchanged.

This expected crosslinking yield needs to be taken into account for any assay development using diazirine lipids to predict experimental yield and necessary biochemical parameters. For microscopy assays, starting bifunctional lipid concentrations should be approximately 20 times higher compared to fluorescence lipid probes to achieve similar signal intensity. To an extent, this can be mitigated by the choice of fluorescent dye used for labelling, but it also highlights the need for optimizing other experimental parameters such as UV light sources, where LEDs should be used instead of arc lamps, and reaction conditions for click chemistry derivatizations. We anticipate that our results will provide a valuable reference point for designing crosslinking studies based on diazirine lipids, which are increasingly used for mapping intracellular lipid localization, measuring lipid transport and identifying lipid-protein interactions.

## Supporting information

Kirschbaum et al 2026 Supplementary Information

## ASSOCIATED CONTENT

Materials and methods, supplementary mass spectra and fragment assignments (PDF). Raw mass spectra are accessible via Figshare (DOI: 10.25446/oxford.30814682).

## AUTHOR INFORMATION

### Author Contributions

The manuscript was written through contributions of all authors. All authors have given approval to the final version of the manuscript.

### Notes

The authors declare no competing financial interest.

## ACKNOWLEDGMENT

The authors acknowledge funding by the Leopoldina fellowship program of the German National Academy of Sciences Leopoldina (LPDS 2023-07; C.K.) and a Wellcome Trust grant (221795/Z/20/Z; C.V.R.). This work was supported by an Allen Distinguished Investigator Award, a Paul G. Allen Frontiers Group advised grant of the Paul G. Allen Family Foundation to A.N.

## Notes

### Competing Interest Statement

The authors have declared no competing interest.

