## Supplementary material for "Bifunctional Lipid-Protein Crosslinking Efficiency and Reaction Products": Kirschbaum et al 2026 Supplementary Information

### Crosslinking Efficiency and Reaction Pathways of Diazirine Lipids: Insights from a Protein Model System

### Supplementary Figures and Tables

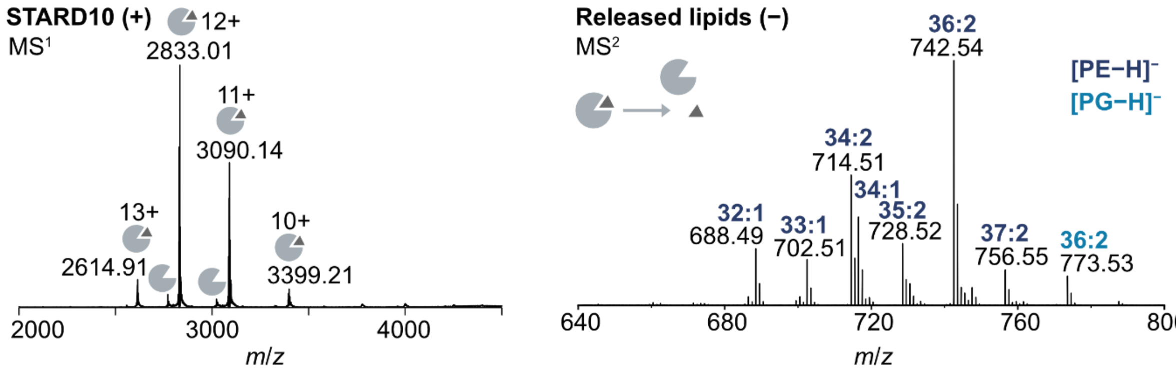

**Figure S1.** Native mass spectrometry of STARD10 expressed in *E. coli*. The native mass spectrum shows that the majority of STARD10 is bound to bacterial phospholipids at a 1:1 ratio. The copurified lipids were identified by MS$^2$ as bacterial phosphatidylethanolamines (PE) and phosphatidylglycerols (PG).

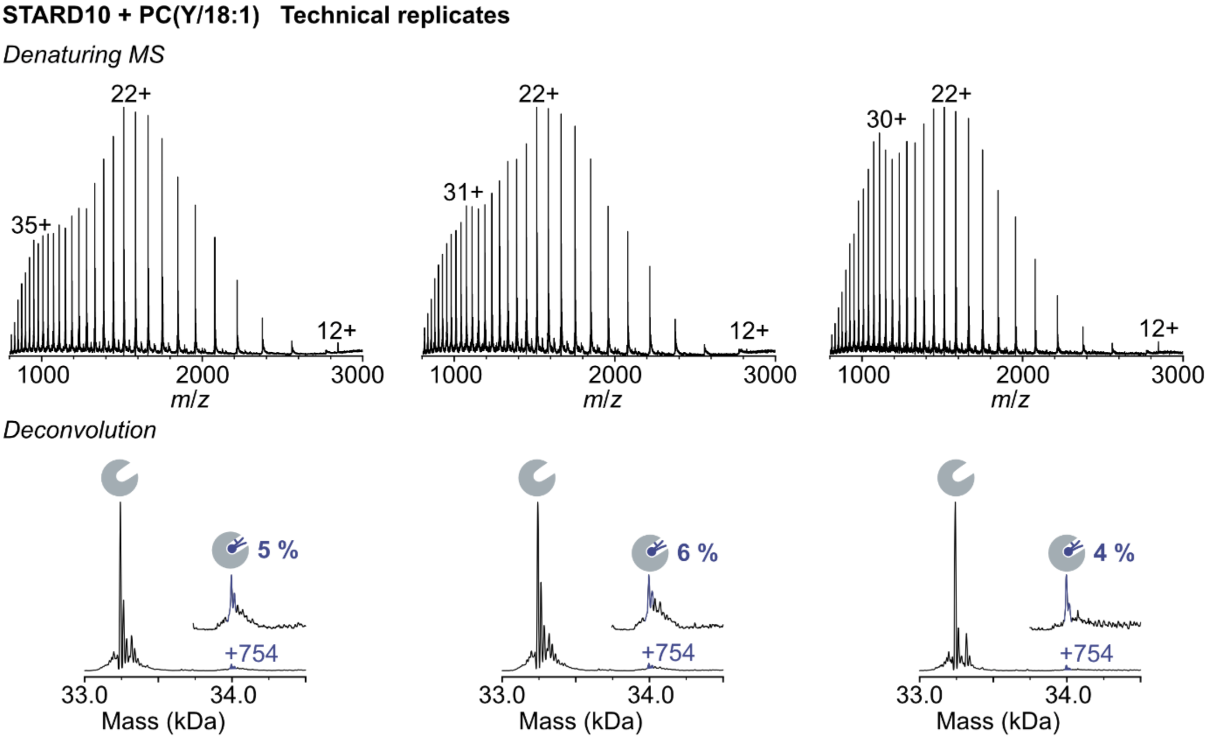

**Figure S2.** Technical replicates: quantification of PC(Y/18:1) crosslinked to STARD10 after UV photoactivation. Deconvolution of denaturing mass spectra reveals crosslinking ratios between 3–5 % (assuming ca. 1 % contribution from non-covalent binding).

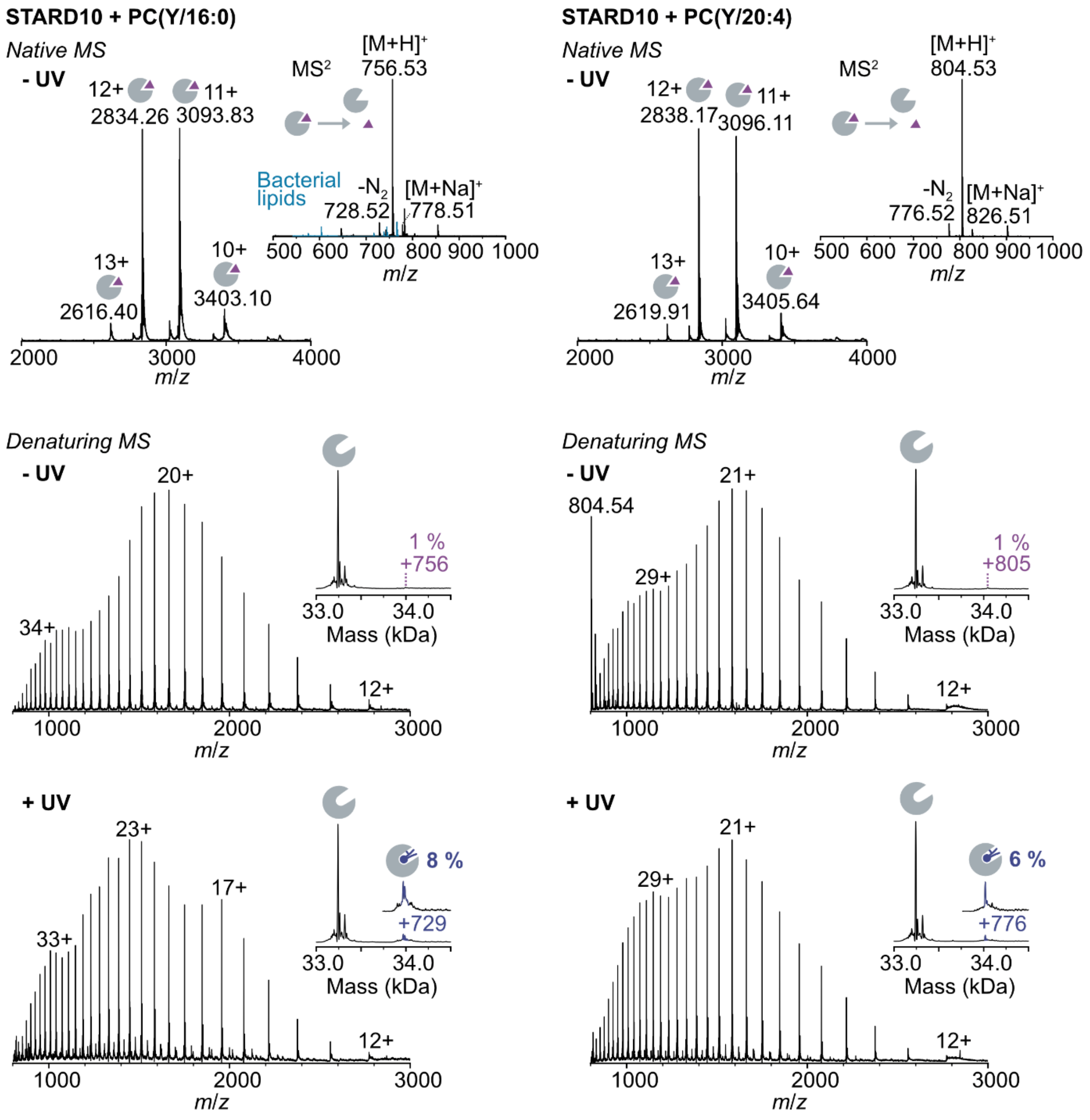

**Figure S3.** Crosslinking between STARD10 and bifunctional PC probes with varying acyl chains (PC(Y/16:0) and PC(Y/20:4)). Native MS confirms binding of bifunctional lipids to STARD10 in a 1:1 stoichiometry. The crosslinking efficiency is determined by deconvolution of denaturing mass spectra.

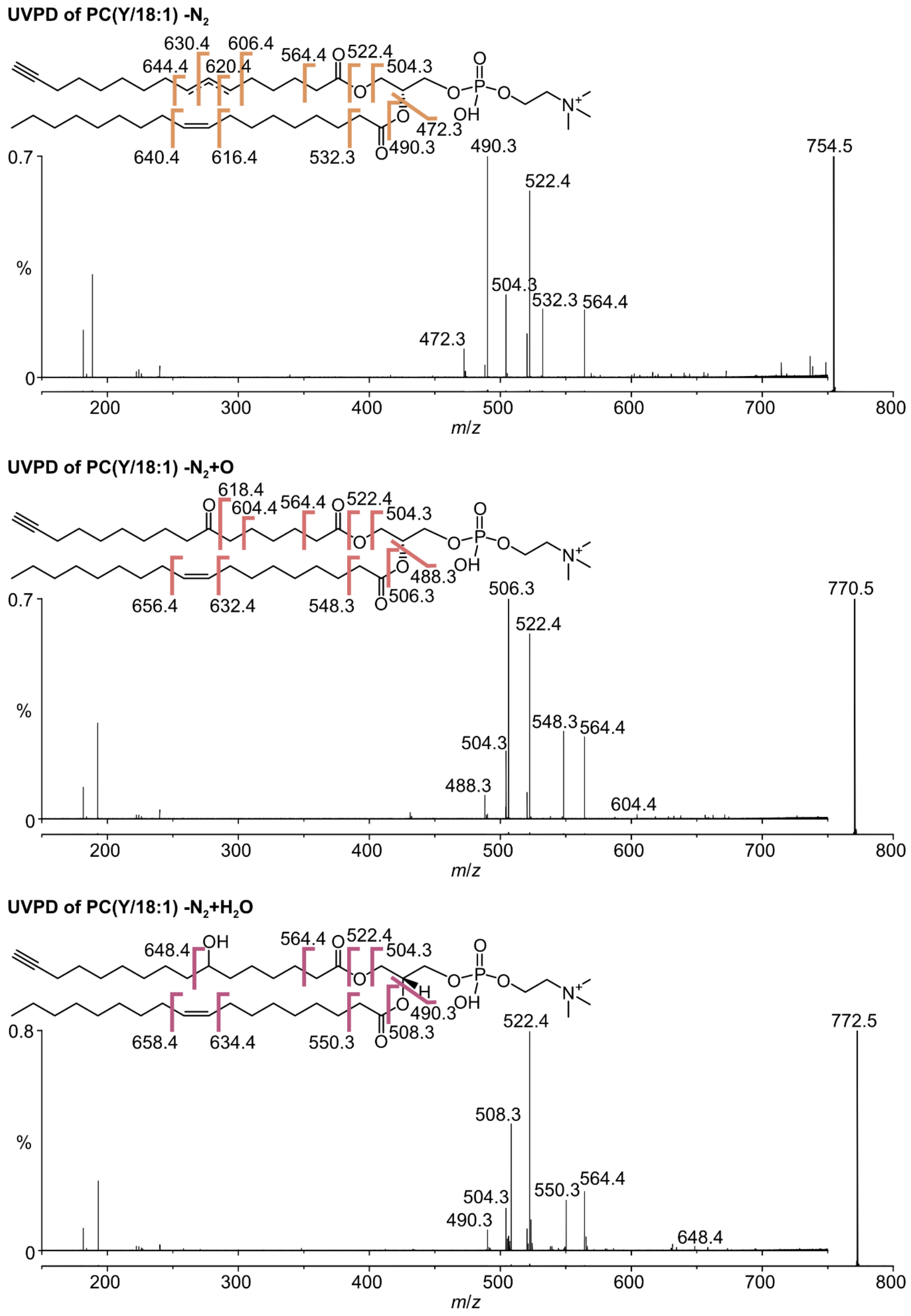

**Figure S4.** Full ultraviolet photodissociation spectra of PC(Y/18:1) photoreaction products with fragments assigned.

**Table S1.** UVPD fragment assignments for PC(Y/18:1) photoreaction products.

| Assignment | Expected mass | Measured mass | Mass error (ppm) |
|---|---|---|---|
| **PC(Y/18:1) -$N_2$** | 754.5381 | 754.5383 | 0.3 |
| Loss of *sn*-1 acyl | 564.3660 | 564.3668 | 1.4 |
| | 522.3554 | 522.3562 | 1.5 |
| | 504.3449 | 504.3456 | 1.4 |
| Loss of *sn*-2 acyl | 532.3034 | 532.3042 | 1.5 |
| | 490.2928 | 490.2936 | 1.6 |
| | 472.2823 | 472.2831 | 1.7 |
| *sn*-1 double bond | 606.4129 | 606.4142 | 2.1 |
| | 630.4129 | 630.4143 | 2.2 |
| | 620.4286 | 620.4288 | 0.3 |
| | 644.4286 | 644.4301 | 2.3 |
| *sn*-2 double bond | 616.3973 | 616.3978 | 0.8 |
| | 640.3973 | 640.3976 | 0.5 |
| | 754.5381 | 754.5383 | 0.3 |
| **PC(Y/18:1) -$N_2$ +O** | 770.5330 | 770.5333 | 0.4 |
| Loss of *sn*-1 acyl | 564.3660 | 564.3668 | 1.4 |
| | 522.3554 | 522.3561 | 1.3 |
| | 504.3449 | 504.3456 | 1.4 |
| Loss of *sn*-2 acyl | 548.2983 | 548.2991 | 1.5 |
| | 506.2877 | 506.2884 | 1.4 |
| | 488.2772 | 488.2779 | 1.4 |
| ketone | 604.3973 | 604.3981 | 1.3 |
| | 618.4129 | 618.4137 | 1.3 |
| *sn*-2 double bond | 632.3922 | 632.3929 | 1.1 |
| | 656.3922 | 656.3926 | 0.6 |
| **PC(Y/18:1) -$N_2$ +$H_2O$** | 772.5487 | 772.5475 | -4.9* |
| Loss of *sn*-1 acyl | 564.3660 | 564.3668 | 1.4 |
| | 522.3554 | 522.3563 | 1.7 |
| | 504.3449 | 504.3457 | 1.6 |
| Loss of *sn*-2 acyl | 550.3139 | 550.3148 | 1.6 |
| | 508.3034 | 508.3041 | 1.4 |
| | 490.2928 | 490.2936 | 1.6 |
| alcohol | 648.4235 | 648.4250 | 2.3 |
| *sn*-2 double bond | 634.4078 | 634.4085 | 1.1 |
| | 658.4078 | 658.4091 | 2.0 |

* overlap with PC Y/18:1 -$N_2$ +O isotope

### Materials and Methods

#### Expression and purification of STARD10

Codon-optimized DNA corresponding to STARD10 with an N-terminal hexahistidine tag, maltose binding protein and TEV protease cleavage site was cloned into a modified pET28b vector using Gibson assembly. After transformation into *E. coli* C43(DE3) cells (New England Biolabs), the bacteria were grown overnight at 37 °C in Luria Broth under kanamycin selection (50 µg/L). Each liter of expression culture was inoculated with 10 mL overnight culture and grown under kanamycin selection at 37 °C until the cell density reached $OD_{600}$ = 0.5. Protein expression was induced by adding IPTG (0.5 mM), and cells were grown for 4 h at 30 °C. Cell pellets were harvested by centrifugation (5000 × g, 15 min).

The bacteria were lyzed by four passes through a microfluidizer (30,000 psi) in 20 mL lysis buffer per liter of culture (50 mM Tris pH 8.0, 300 mM NaCl, 2.5 mM BME, 1 protease inhibitor tablet per 50 mL). The lysate was cleared by centrifugation (20,000 × g, 20 min), filtered through a 0.45 µm syringe filter and loaded onto a 5-mL Ni-NTA column. The column was washed with 25 mL wash buffer (20 mM Tris pH 8.0, 300 mM NaCl, 20 mM imidazole, 2.5 mM BME) before and with 50 mL wash buffer after loading. The protein was eluted with 25 mL elution buffer (20 mM Tris pH 8.0, 300 mM NaCl, 300 mM imidazole, 2.5 mM BME). After dialysis with TEV protease overnight against imidazole-free buffer, STARD10 was separated from His-tagged maltose binding protein by reverse Ni-NTA affinity column chromatography. The protein was concentrated using a spin filter and further purified on a Superdex 200 10/300 Increase gel filtration column (20 mM Tris pH 8.0, 100 mM NaCl, 2.5 mM BME).

#### Synthesis of bifunctional lipids

The bifunctional lipid probes PC(Y/16:0), PC(Y/18:1) and PC(Y/20:4) were synthesized as reported previously.[1]

#### Preparation of liposomes

Liposomes containing 80 % bifunctional PC and 20 % cholesterol were prepared in PBS buffer. Briefly, cholesterol in chloroform (14.5 µL, 10 mg/mL) was mixed with bifunctional PC in chloroform (150 µL, 10 mM) in a glass tube. The lipid film was dried under an argon flow and resuspended in 500 µL PBS, resulting in a final PC concentration of 3 mM. Liposomes were extruded through a mini extruder (Avanti Research) using 100 nm polycarbonate membranes and were stored at 4 °C.

#### Protein–lipid crosslinking

Protein–lipid crosslinking was performed in a 96-well plate. STARD10 (20 µM) was incubated with liposomes (15 µL) in a total volume of 80 µL PBS for 10 min at room temperature. Crosslinking was performed by irradiation with high-powered 365 nm LEDs (Violumas) for 3 s. Control samples were not irradiated. P-6 biospin columns (Bio-Rad Laboratories) were used to remove excess lipids and change the buffer to ammonium acetate (200 mM, pH 7). Samples were flash-frozen in ammonium acetate and stored at −80 °C until analysis.

#### Native mass spectrometry

Native mass spectra were recorded on an Orbitrap Eclipse Tribrid mass spectrometer (Thermo Fisher Scientific). Samples were ionized by nano electrospray ionization using borosilicate capillaries pulled in-house with a P97 Micropipette Puller (Sutter Instrument Corporation) and gold-coated with an AgarAuto Sputter Coater. Native mass spectra were obtained in Intact Protein mode using the Orbitrap mass analyzer (typically *m*/*z* 1000–6000, resolution = 15,000

@ *m*/*z* 200). To identify bound lipids, protein-lipid complexes were isolated in the ion trap using a wide isolation window (100 Da) and activated by higher-energy collisional dissociation (HCD; 12 % NCE). Spectra of released lipids were recorded in the Orbitrap at high resolution (*m*/*z* 500–1000, resolution = 500,000 @ *m*/*z* 200).

**Denaturing mass spectrometry**

For denaturing MS, samples in ammonium acetate buffer were diluted 1:1 with a 9:1 acetonitrile:isopropanol mixture containing 2% formic acid, resulting in a mixture of ca. 50% water, 45% acetonitrile and 5% isopropanol containing 100 mM ammonium acetate and 1% formic acid. Mass spectra were obtained on an Orbitrap Eclipse Tribrid mass spectrometer in Small Molecule mode using the Orbitrap mass analyzer (typically *m*/*z* 500–4000, resolution = 15,000 @ m/z 200). For quantification of protein-lipid crosslinking products, mass spectra were deconvolved using UniDec (*m*/*z* 800–3000, background subtraction, charge range 1–50, mass range 30,000–40,000 Da, 1.0 Da mass sampling, 0.1 FWHM), and the resulting peak areas were integrated (±10 Da).

**Ultraviolet photodissociation**

Ultraviolet photodissociation (UVPD) spectra of lipids in the denatured protein samples were recorded on an Orbitrap Eclipse Tribrid mass spectrometer equipped with a 213 nm UV laser. The lipids were isolated using the quadrupole using a narrow isolation width (1.5 Da) to select monoisotopes. UVPD spectra were recorded in the Orbitrap at high resolution (*m*/*z* 150–1000, resolution 240,000 @ *m*/*z* 200) with an irradiation time of 1000 ms. Spectra were averaged over at least 30 individual scans.
